# The small membrane protein YohP induces membrane depolarization and ppGpp accumulation in *Escherichia coli*

**DOI:** 10.1101/2025.05.06.652344

**Authors:** Ana Natriashvili, Nahid Mohammadsadeghi, Martin Milanov, Eva Smudde, Isabel Prucker, Henning J. Jessen, Iulia Carabadjac, Heiko Heerklotz, Pedro H.C. Franco, Julian D. Langer, Hans-Georg Koch

## Abstract

Small membrane proteins represent an abundant and ubiquitous class of proteins that are often up-regulated when cells encounter unfavorable conditions, yet details about their exact function are largely missing. In bacteria, these proteins consist of typically less than 50 amino acids and contain a single transmembrane domain, but lack any detectable catalytic activity. Thus, the benefit of producing these proteins during stress conditions is unknown. In the current study we used a multidisciplinary approach to determine the function of the 27 amino acid long protein YohP in *E. coli*. Our proteomics approach revealed that YohP production leads to an up-regulation of proteins involved in membrane protection and to a down-regulation of many enzymes involved in key metabolic processes, such as nucleotide biosynthesis. Further biochemical characterizations revealed increased cardiolipin content in the membrane, a partial dissipation of the membrane potential and reduced membrane fluidity in YohP-containing membranes. Finally, our data show that YohP production induces the stringent response and leads to elevated levels of (p)ppGpp. Overall, our data indicate that the YohP-induced proteome and membrane changes initiate a state of metabolic silencing that protects *E. coli* against stress and helps to conserve cellular resources.

## Introduction

Bacteria are facing constant changes in their environment and the ability to execute counteracting strategies is therefore of utmost importance for their survival. This multilayered response includes the well-established transcriptional regulation via transcription factors and regulatory RNAs (1–4), translational control (5–7) and the production of small stress-responsive signaling molecules, such as the hyperphosphorylated guanine nucleotides pppGpp and ppGpp (8,9). The synthesis of small proteins of typically less than 50 amino acids serves as an additional bacterial stress-response strategy and this applies in particular to small membrane proteins (SMPs). These proteins consist of a single transmembrane domain with only a few amino acids residing in the cytoplasm or periplasm (10–14). Due to their small size and their predicted simple topology, SMPs are unlikely to act catalytically, but instead often execute their function by interacting with other membrane proteins. This has been shown for the small membrane proteins MgtS and MgtU, which protect the magnesium transporters MgtA and MgtB from degradation (15). Another example is AcrZ, which stimulates export of antibiotics via the multi-drug efflux pump AcrAB (16) or MgrB, which inhibits the sensor kinase PhoQ and prevents PhoQP hyperactivation (17). SMPs are also important for maintaining the energy status of bacterial cells by interacting with terminal oxidases (18–24).

In addition to these few well characterized examples, several hundreds of uncharacterized soluble and membrane-bound small proteins are expected to exist in *E. coli* (13,14), for which a functional characterization is still missing. This also still applies to many of the experimentally verified small proteins (25,26). Examples are YncL, which consists of just 31 amino acids and is up-regulated during heat-shock in *E. coli*, or YohP, which consists of 27 amino acids and shows increased levels when cells were exposed to hydrogen peroxide or SDS+EDTA treatment (27). YohP also serves as a model protein for studying the membrane insertion pathway of SMPs. This pathway involves a post-translational recognition by the signal recognition particle (SRP) and a subsequent targeting to the SecYEG translocon or, alternatively, to the YidC insertase (28,29). Under stress conditions, *e.g.* when the SRP pathway is inhibited by accumulating (p)ppGpp (30), YohP insertion occurs SRP-independently via mRNA targeting (31).

YohP is localized to the inner membrane with a mainly N_out_-C_in_ topology (29), but it might also acquire a dual topology, according to GFP- and PhoA-reporter fusion studies (10). YohP shows a strong propensity for dimerization via an unusual glycine motif (29) and high YohP levels cause nucleoid condensation. Nucleoid condensation via nucleoid-associated proteins is an important factor of global gene regulation in response to environmental changes. DNA-interacting proteins have only restricted access to a condensed nucleoid, which reduces gene expression and subsequently leads to decreased metabolic activity, which in turn can protect cells against unfavorable conditions (32). This is in line with data showing that nucleoid condensation is also observed upon induction of some membrane-permeabilizing toxins or upon contact with antimicrobial peptides (33–36). SMPs share some common features with membrane-acting toxins of toxin-antitoxin systems or membrane-targeting antimicrobial peptides, although they don’t show strong sequence conservation on the amino acid level (33,37–40). Besides their small size of usually less than 50 amino acids, they all consist of a single α-helical transmembrane domain with the C-terminus predicted to reside mainly in the cytosol. In addition, their transmembrane domains are often flanked by positively charged residues, which are important for membrane interaction. This raises the possibility that some SMPs act like antimicrobial peptides or membrane-acting toxins by disrupting the membrane integrity.

In the current study, we determined the consequences of YohP production in *E. coli* by highly complementary proteomic, lipidomic and biochemical approaches and our data demonstrate that YohP causes a non-lethal reduction in membrane potential, which induces a significant proteome rearrangement and stimulates the stringent response.

## Results

### YohP synthesis significantly alters the *E. coli* proteome

YohP is an inner membrane protein of just 27 amino acids that form a single transmembrane helix, which is mainly oriented in a N_out_-C_in_ topology (29,31). For monitoring growth-phase dependent YohP synthesis, we used an *E. coli* MG1665 variant that contained a C-terminally SPA-tagged *yohP* copy in the chromosome under its native promoter (MG1665-*yohP*-SPA) (12). In addition, we expressed in MG1665 a plasmid-encoded *yohP* construct under the *T7*-dependent promoter in plasmid pRS1 (29). The plasmid-encoded YohP construct contained a triple-Flag tag at the C-terminus and expression was induced with 1mM IPTG. Both *E. coli* strains were grown in LB medium and analyzed for YohP-levels via western-blotting using antibodies against the Flag-tag (**Fig. 1A**). This revealed a growth-phase dependent increase of the native YohP levels, starting at late exponential/mid stationary phase (OD 1.5 – 2.5) and increasing up to late stationary phase (OD 4.0). YohP produced from the plasmid-encoded *yohP* copy was already detectable at early exponential phase (OD 0.5) and increased only moderately up to stationary phase (OD 4.0). The SPA-tag consist of an 8 kDa calmodulin-binding peptide in addition to the triple-Flag tag and therefore the chromosomally tagged YohP version migrates above the plasmid-encoded YohP version. Immune detection of the inner membrane protein YidC served as loading control. In summary, these data demonstrate that native YohP is produced when *E. coli* cells enter stationary phase.

**Fig. 1.**
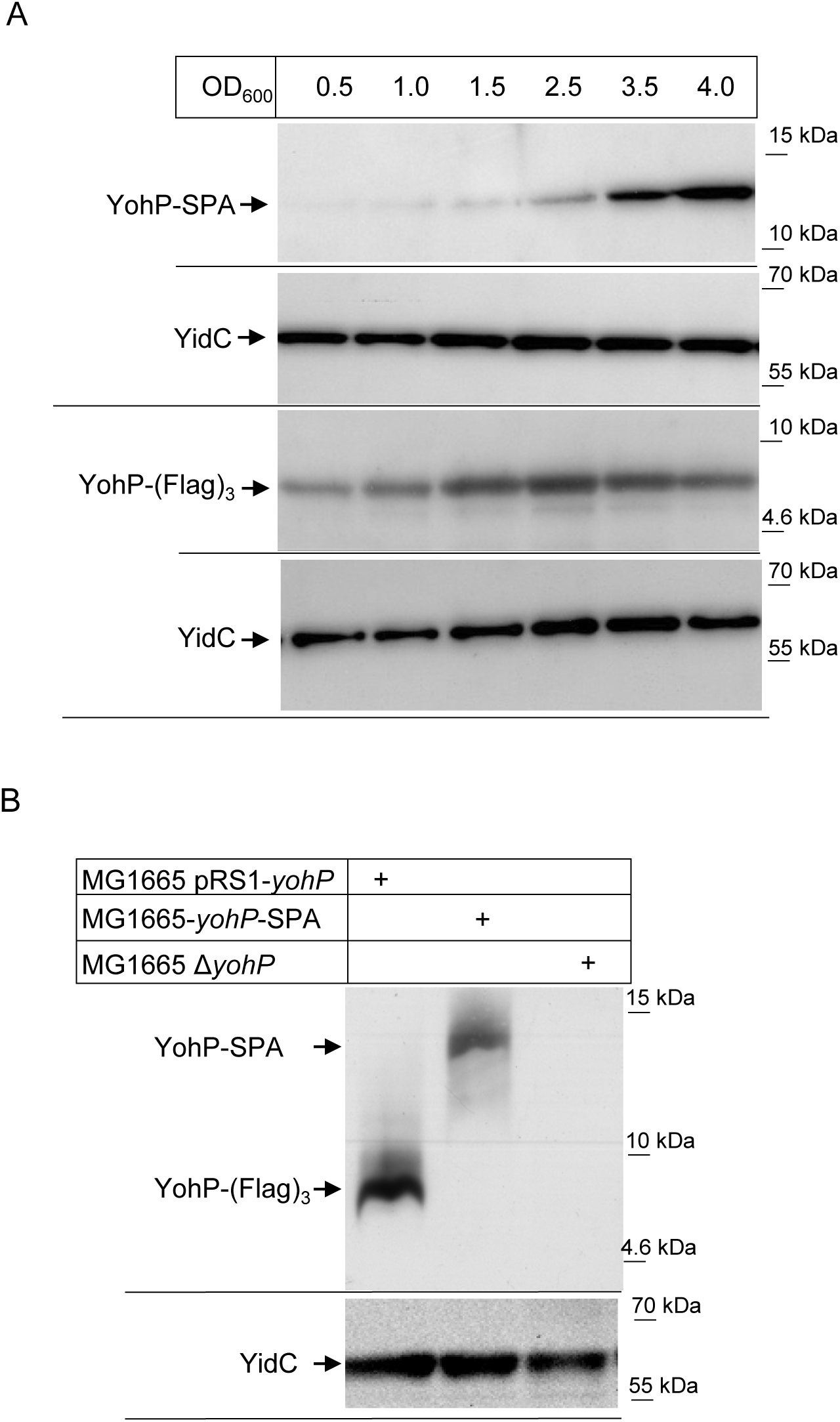
Native YohP increase when cells enter stationary phase. **A**. *E. coli* MG1665 containing a chromosomally SPA-tagged YohP under its native promoter (YohP-SPA) and *E. coli* containing a plasmid-containing Flag-tagged YohP copy (YohP-Flag_3_) were grown on LB medium up to different ODs and 2 x 10^8^ cells were precipitated with 5% trichloroacetic acid (TCA). After centrifugation, samples were denatured and separated on 16.5 % Tris-Tricine SDS-PAGE followed by western transfer and immune detection with α-Flag antibodies. The expression of the plasmid-encoded *yohP* copy was induced at OD 0.4 with 1 mM IPTG. Antibodies against YidC served as a loading control. **B**. Strains were grown on phosphate-buffered rich medium and the strain containing pRS1-*yohP* was induced at OD 0.2 with 1 mM IPTG for 2 hours. Samples were then processed as in **A**. Antibodies against the inner membrane protein YidC served as loading control. Shown are representative blots of at least 3 independent biological replicates.

YohP production has been linked to stress conditions (12,26,27), but its function is still unknown. We therefore performed mass-spectrometry (MS) based proteomics, comparing wild type cells with a Δ*yohP* strain and a wild type strain carrying the plasmid-encoded His-tagged pRS1-*yohP* copy. The MG1665 Δ*yohP* strain was constructed via λ-red recombination (41), and the absence of YohP did not cause any growth defect on LB-medium (**Fig. S1A**). However, when the expression conditions of the pRS1-*yohP* copy were adjusted to levels corresponding to the native levels that were observed in the MG1665-*yohP*-SPA strain during stationary phase (**Fig. 1B**), we noticed a reduced growth rate (**Fig. S1A**). This indicates that *E. coli* growth is inhibited when significant amounts of YohP are already produced during the exponential phase.

To investigate the proteome remodeling deriving from *yohP* deletion or *yohP* induction, in comparison to wild type *E. coli*, we conducted MS-based proteomics in a data-independent acquisition mode (n = 3 per group). We chose to monitor the effects in exponentially grown cells to exclude stationary phase effects, such as the induction of the RpoS-response (42,43). With our workflow, we quantified more than 2,500 proteins per sample (**Fig. 2A**), achieving an average coefficient of variation (CV) of approx. 10% at the protein level (**Fig. 2B**). This highly comprehensive dataset provided a solid foundation for a robust comparison of the *yohP* deletion and *yohP* induction strains against wild type *E. coli*. Based on clustering analysis, we observed a high degree of similarity among all sample groups (R > 0.95) **(Fig. 2C**), with the *yohP* induction strain showing a more prominent difference. We then carried out differential expression analysis using the DEqMS algorithm (44), implemented in the MS-DAP-R package (45), which accounts for the dependency of variance on the number of peptides used for quantification. With the ability to quantify over 30,000 peptides per sample, our workflow enabled precise identification of significantly upregulated and downregulated proteins in the analyzed strains.

**Fig. 2.**
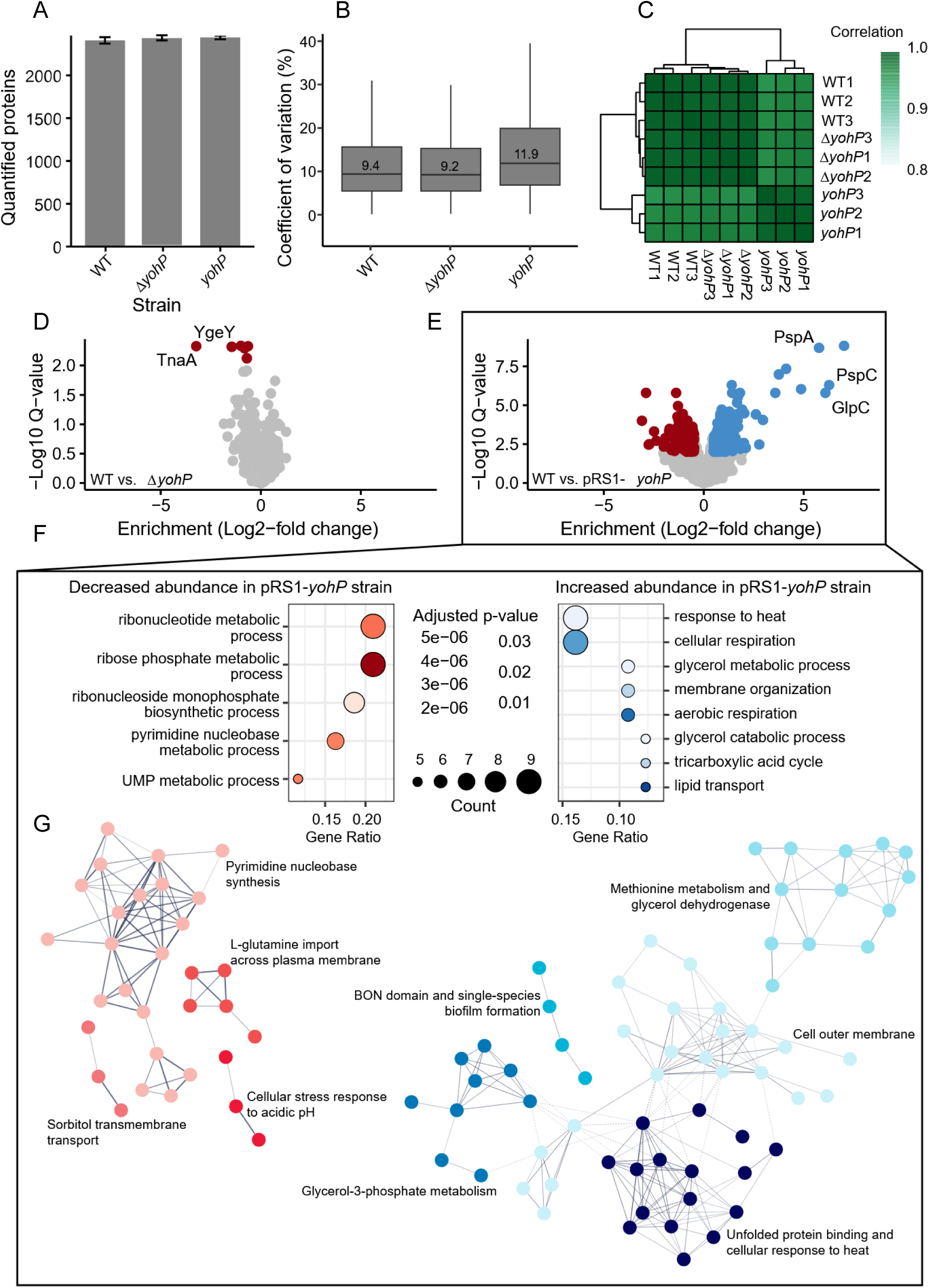
High quality MS-data provides insights into the role of YohP. **A**. Average number of quantified proteins in each measured sample group (n = 3). Error bars are represented in terms of standard deviation. **B**. Boxplot representation of the average coefficients of variation (%) for quantified proteins in each sample group (n =3). **C**. Correlation heatmap and clustering analysis of the measured samples. Correlation values in the legend are in terms of coefficient of correlation (R). **D**. Volcano plot for the comparison between wild-type *E. coli* and the *yohP*-deletion strain. Significantly differentially abundant proteins (q-value < 0.05 and fold-change > 2) are highlighted in red for decreased abundance in the mutant strain. **E**. Volcano plot for the comparison between wild-type *E. coli* and the *yohP*-induction strain. Significantly differentially abundant proteins (q-value < 0.05 and fold-change > 2) are highlighted in red for decreased abundance in the mutant strain, and in blue for increased abundance. **F**. Dotplots for the GO classification enrichment analysis for the comparison between wild-type *E. coli* and the *yohP*-induction strain. GO classifications for Biological Processes (BP) were selected. **G**. String networks for the comparison between wild-type *E. coli* and the *yohP*-induction strain. Only differentially abundant proteins (q-value < 0.05 and fold-change > 2) were selected for the network analysis, and non-interacting nodes were removed. K-means clustering analysis of the network led to significant nodes (p < 0.05) grouped by different colors.

The comparison of the proteomes of the Δ*yohP* strain and the wild-type *E. coli* showed only a small number of proteins with decreased abundance in the absence of YohP and no protein with increased abundance (**Fig. 2D**; **Table S1**). The strongest effect was observed for tryptophanase (TnaA), which is required for the production of the signaling molecule indole. The putative peptidase YgeY was also down-regulated. YgeY was identified in a screen for envelope biogenesis mutants (46), but so far, no clear functions have been assigned. Small effects were also observed for maltose transport proteins MalK and MalE, the ketol-acid reduktoisomerase IlvC and the L-asparaginase AnsB. A recent report demonstrates that MalK is down-regulated when the SOS DNA-damage response is induced (47). However, we did not see any growth impairment of the Δ*yohP* strain on maltose-McConkey-agar (**Fig. S1B**), indicating that the Δ*yohP* strain can still utilize maltose as carbon source.

In contrast to the Δ*yohP* strain, the YohP-producing strain showed 80 proteins that were significantly upregulated (**Fig. 2E**; **Table S2**) and 52 down-regulated proteins (**Fig. 2E**; **Table S2**). Based on gene ontology (GO) classification **(Fig. 2F**), the most strongly upregulated proteins corresponded to heat-shock proteins/chaperones, proteins involved cellular and aerobic respiration, in membrane organization and lipid transport, as well as glycerol-3-phosphate metabolism. The strongest upregulations were observed for the phage-shock proteins PspA and PspC and for the anaerobic glycerol-phosphate dehydrogenase (GlpABCD). PspA, PspC and GlpC have been shown to respond to membrane damage, induced *e.g.* by organic solvents or upon dissipation of the membrane potential (48,49). GO classification for cellular compartment revealed significant enrichments for proteins localized to the cell envelope and outer membrane (**Fig. S2**).

Many proteins involved in nucleotide biosynthesis were downregulated upon YohP production (**Fig. 2F**). This included carbamoylphosphate synthase (CarB), which initiates pyrimidine biosynthesis, aspartate transcarbamylase (PyrB), which catalyzes the second step of pyrmidine biosynthesis and many more proteins involved in pyrimidine synthesis (PyrD, PyrC, Upp). In addition, the uracil transporter UraA, the cytosine transporter CodB and the glutamine transporter GlnHPQ were downregulated. GlnHPQ constitute an ABC-transporter required for high-affinity uptake of glutamine (50), which serves as substrate for the CarB reaction. Some proteins involved in purine metabolism, such as GMP-reductase GuaC, or xanthine-guanine phosphoribosyltransferase Gpt were also down-regulated. In summary, these down-regulations most likely reduce the *de novo* synthesis and salvage pathways of pyrimidine nucleotides, and, to a lesser extent, purine nucleotides.

The production of YohP also reduced the levels of some regulatory proteins, such as AppY, GadE, YfeC or GlnG (NtrC). AppY regulates genes of the anaerobic energy metabolism, such as the hydrogenase-I (*hyaABCDEF*), which we also found to be down-regulated. Furthermore, both AppY and GadE control genes involved in acid resistance (51,52) and both have been linked to the SOS DNA damage control (47). YfeC acts as a dual regulator that is involved in regulating DNA replication, translation and cell envelope biogenesis (53). Finally, NtrBC constitutes a two-component system that controls the transcriptional response to nitrogen starvation and regulates amino acid metabolism (54).

We have also observed a high-degree of interaction between the differentially expressed proteins. Network analysis using STRING (55) revealed that the protein-protein interaction (PPI) enrichment p-value for the downregulated proteins in the YohP-overexpressing strain was smaller than 1e^-16^. The same degree of PPI enrichment was observed for the upregulated proteins. This means that the degree of interaction of the up- and downregulated proteins is significantly higher than what would be expected for a random set of proteins, indicating that they are biologically connected (**Fig. 2G**). We have also verified by k-means cluster analysis similar biological function enrichments to what we have observed in GO-terms analysis **(Fig. 2G**, **Figs. S3 & S4**). Furthermore, we investigated the interactions between up- and downregulated proteins and observed similar PPI enrichment p-values (Fig. S4). These data show that there are interactions between proteins differentially regulated in the YohP-producing strain. Examples include GadE (downregulated), YdeP and AdiY (upregulated), involved in acid resistance, GalE (downregulatd) and CpsG (upregulated), involved in galactose metabolism, and GltS (upregulated) and GlnQ (downregulated), involved in glutamine/glutamate transport.

In conclusion, the deletion of *yohP* has only a minor effect on the *E. coli* proteome and mainly reduces the amount of tryptophanase. In support of this, no detectable phenotype has been observed so far for the Δ*yohP* strain. In contrast, YohP overproduction causes a multi-layered change in the protein composition, with the most striking effects on proteins involved in acid response, membrane damage and nucleotide biosynthesis. This is also reflected by the reduced growth rate of the *yohP*-producing strain (**Fig. S1A**)

### YohP has only a small effect on indole production and does not influence growth of *E. coli* at different pH-values

For a phenotypic validation of the proteomic changes, indole levels were determined in the Δ*yohP* strain and compared to indole levels of the wild type and a Δ*tnaA* strain. Indole levels were initially monitored in cells grown on phosphate-rich medium (INV medium), which was also used for the proteomics approach. However, under those conditions indole was not detectable. We therefore switched to tryptone broth, which has previously been used for monitoring indole levels. Indole was detectable in wild type cells, but not in the Δ*tnaA* strain (**Fig. 3A**). Indole formation in wild type cells reached a maximum at late exponential phase and then declined. In the absence of YohP, the indole production was reduced but not completely diminished (**Fig. 3A**), which is in line with the reduced TnaA levels shown in the proteome analysis of the Δ*yohP* strain. We also monitored the TnaA levels by immune detection and observed slightly reduced but not completely diminished TnaA levels in the Δ*yohP* strain (**Fig. 3B**), supporting the reduced but still detectable indole production in this strain.

**Fig. 3.**
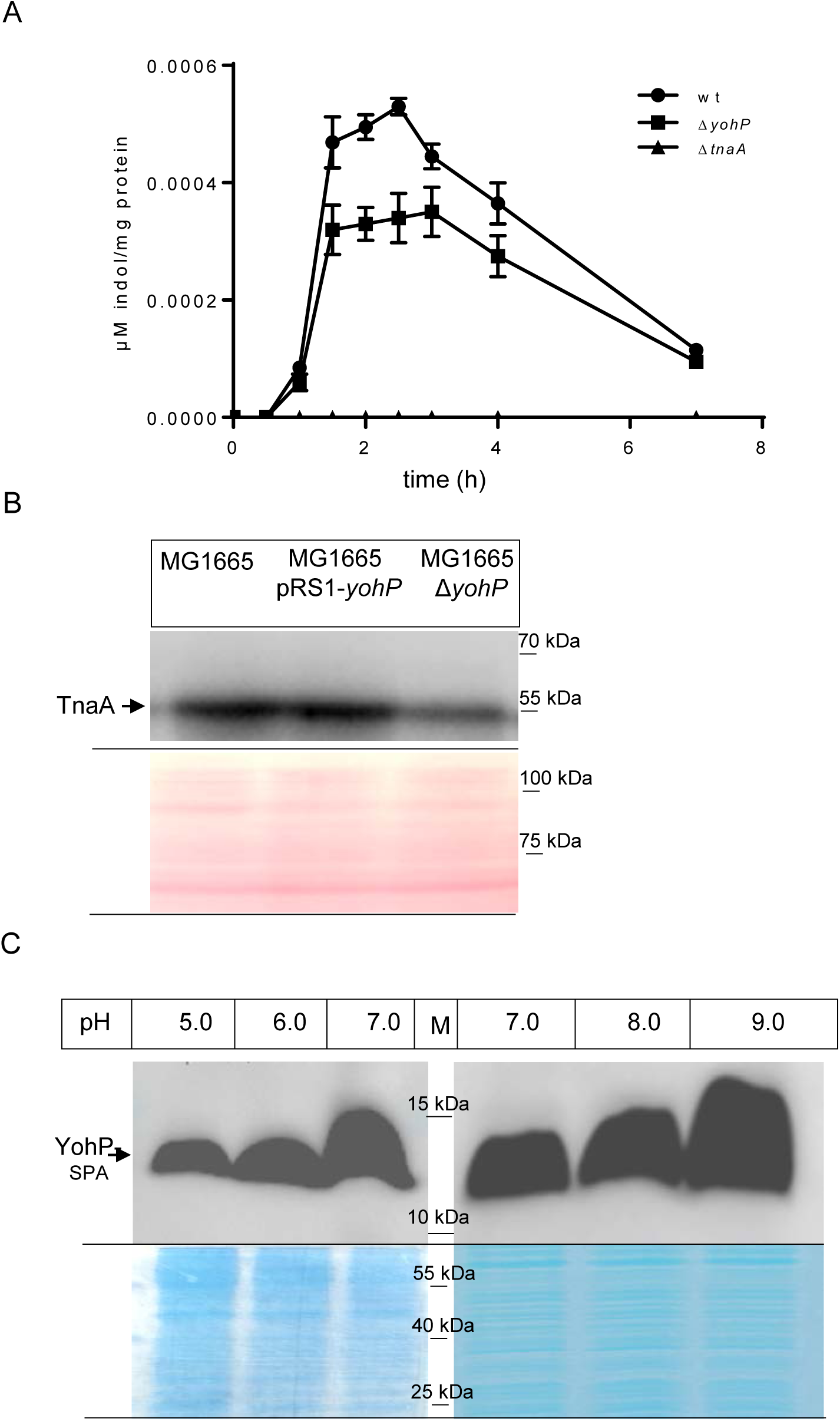
Indole formation is reduced in the absence of YohP and *yohP* expression is increased at basic pHs. **A**. *E. coli* cells were grown on tryptone broth and the secreted indole in the supernatants was quantified after different times of incubation using the Kovacs reagent. Shown are the mean values (n ≥ 2) and the error bars represent the standard deviation. **B**. *E. coli* cells were grown as in **A** up to OD 0.8-1.0 and 2 x 10^8^ cells were processed as in Figure 1 and further analyzed with α-TnaA antibodies. The lower panel displays the Ponceau Red stained membrane used for immune detection. **C**. *E. coli* cells were grown on LB medium adjusted to different pH values up to OD 0.8-1.0 and 2 x 10^8^ cells were processed as in Figure 1. The lower panel corresponds to a Coomassie-stained part of the gel as loading control.

Indole production has been linked to acid resistance in *E. coli*, although there are conflicting results as to whether it inhibits (56) or stimulates (57) the acid resistance system. A possible link between the YohP-levels and acid resistance is also supported by the down-regulation of the transcriptional regulators AppY, and GadE, which are involved in acid resistance (51,52) and of glutamine ABC transporter GlnHPQ, which also influences acid resistance (58). This was analyzed by monitoring YohP expression and cell growth at different pH values. The strain MG1665-*yohP-spa*, containing the chromosomally tagged *yohP*-copy, was grown in M63 medium at different pH-values and the YohP-levels were determined by western blotting using α-Flag antibodies. This revealed that increasing the pH from 5.0 to 9.0 caused an increase in the YohP levels (**Fig. 3C**), demonstrating that YohP-expression is pH-regulated. However, we did not observe any specific growth defect of the Δ*yohP* or *yohP*-overproducing strains at different pH values (**Fig. S1**), suggesting that YohP is not a major factor for controlling *E. coli* growth at different pH values.

### YohP influences membrane composition and fluidity

The up-regulation of PspC, PspA and GlpC in the MG1665 pRS1-*yohP* strain is indicative for membrane damage, and we confirmed the up-regulation of PspC and PspA via western blotting (**Fig. 4A**). The levels of the major stress-responsive σ-factor RpoS were also analyzed for excluding secondary effects, but there were no significant differences between the strains.

**Fig. 4.**
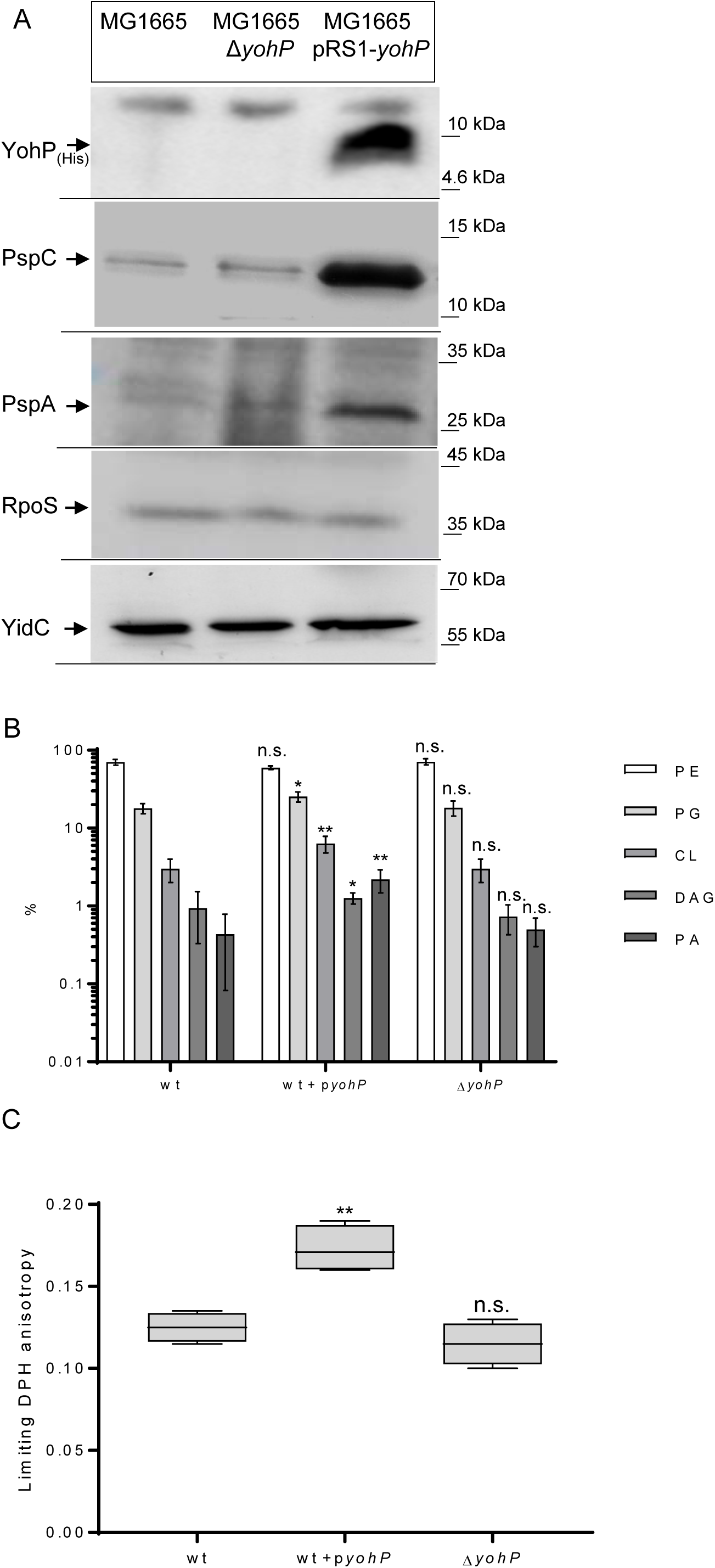
YohP causes membrane damage. **A**. *E. coli* cells were grown on LB medium up to an OD of 0.8 and 2 x 10^8^ cells were processed as in Figure 1 and the membrane was decorated with the indicated polyclonal antibodies, or, in case of YohP, with monoclonal α-His antibodies. **B**. Lipidomic analyses of isolated inner membrane vesicles (INVs) derived from the indicated strains. The extraction, analysis and quantification was performed by Lipotype GmbH, Dresden, Germany, as described in Material and Methods. Shown are the percentage of the main *E. coli* lipids (phosphatidylethanolamine, PE; phosphatidylglycerol, PG, cardiolipin, CL, diacylglycerol DAG and phosphatidic acid, PA). The error bars reflect the standard deviation (n = 3). **C**. Membrane order in INVs of the indicated strains (1mg protein/ml) was monitored in terms of the limiting fluorescence anisotropy of the probe 1.6-Diphenyl-1,3,5-hexatriene (DPH), *i.e.*, the value approached by the anisotropy decay for infinite time after excitation. Shown are the results of three independent biological replicates with three technical replicates each. To determine the significance of the results, the P-value was calculated using the ‘unpaired, two-tailed t-test’ of the program Graph Pad PRISM 6 or the one-way Anova analyses and Turkey’s honest significance test using the values in the wild type strain as reference. The P-values are depicted as asterisks (*) above the graphs as following: n.s. = P > 0,05; * = P ≤ 0,05; ** = P ≤ 0,01.

Next, the influence of YohP production on the lipid composition of the *E. coli* membrane was determined by a lipidomics approach using sucrose-gradient purified inner membrane vesicles (INVs). The most striking difference was a more than two-fold increase of the cardiolipin content in the pRS1-YohP containing strain. The diacylglycerol (DAG) and phosphatidic acid (PA) contents also increased in this strain, but these lipids generally accounted for less than 2% of the total lipid content (59). In the Δ*yohP* strain, the lipid content was comparable to the wild type (**Fig. 4B**).

Both DAG and cardiolipin influence membrane stability (60). DAG has been suggested to seal membranes (61), while an increase in cardiolipin is observed when the ΔpH is reduced due to uncouplers (62). For monitoring the effect of YohP on the physicochemical properties of membranes, fluorescence anisotropy using the fluorescent probe diphenylhexatriene (DPH) (63) was employed. In comparison to wild type INVs, INVs from the YohP-producing strain showed increased limiting anisotropy, indicative for increased lipid order and reduced membrane fluidity **(Fig. 4C)**. The rotational correlation times of DPH did not differ significantly (not shown). In contrast, INVs of the Δ*yohP* strain showed values comparable to wild type INV. Considering that cardiolipin is expected to increase membrane fluidity (60), the increased rigidity of the membrane is likely a direct effect of the presence of YohP in the membrane rather than an indirect effect via changes in the lipid composition.

### The proton motive force is reduced in the presence of YohP

For monitoring the effect of YohP on the membrane potential, cells were stained with the slow-responsive, potential-sensitive dye DiBAC_4_(3) (Bis-(1,3-Dibutylbarbituric Acid) Trimethine Oxonol). DiBAC_4_(3) enters depolarized cells and binds to intracellular proteins and to membranes. Increased depolarization results in increased influx of the anionic dye and increases fluorescence. Thus, DiBAC_4_(3) fluorescence at 530 nm after excitation at 485 nm is proportional to the membrane depolarization. This was confirmed by treating *E. coli* wild type cells with 5 mM of the uncoupler CCCP (cyanide-*m*-chlorophenyl hydrazine). In CCCP-treated cells, a strong increase in DiBAC_4_(3) fluorescence was observed (**Fig. 5A**). Increased fluorescence was also observed in *yohP*-expressing cells, while the Δ*yohP* strain showed a fluorescence comparable to the wild type. As an additional control, we also analyzed YqjD-producing cells. YqjD is a small C-tail anchored membrane protein that like YohP contains a single transmembrane domain and also a high propensity for dimer formation (64). However, YqjD-producing cells did not show increased fluorescence, demonstrating that YohP insertion into the *E. coli* membrane specifically impairs the proton motive force (pmf). Cells expressing YohP were also more sensitive towards CCCP treatment (**Fig. 5B**). This was monitored by determining the number of viable cells after treatment with increasing CCCP concentrations.

**Fig. 5.**
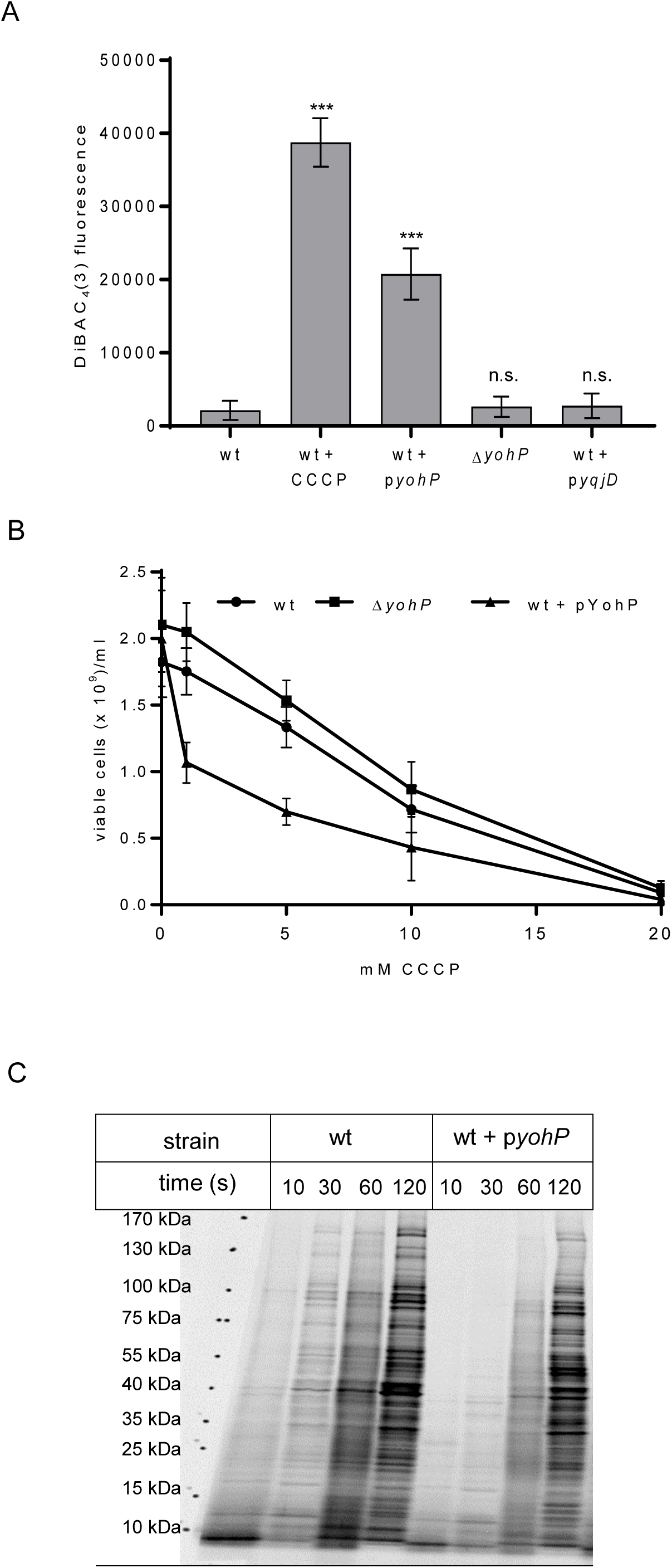
YohP dissipates the membrane potential. **A**. The indicated strains were grown on LB medium and YohP production from the plasmid was induced at OD 0.5 with 0.2 % arabinose, followed by further growth at 37 °C for 2 h. Aliquots of 8 x 10^8^ cells were collected from the samples, washed once with sterile PBS and further incubated for 30 min at 37 °C with either PBS or 5 mM CCCP (Carbonyl-cyanide-m-chlorophenyl hydrazine, dissolved in DMSO). Samples were further processed as described in Material and Methods. DiBAC4(3) fluorescence emission at 530 nm was measured in 96 well plates after excitation at 485 nm. Statistical analyses were performed on three independent biological replicates as described in Fig. 4. **B**. Cells were grown as in **A** and treated with different CCCP concentrations or with DMSO as a control for 1 h at 37 °C. The number of viable cells was then counted using the QUANTOM TxTM Microbial Cell Counter and the QUANTOM^TM^ Viable Cell Staining Kit as described in Material and Methods. Shown are the mean values of three independent biological replicates with two technical replicates each (n = 6). **C**. Global protein synthesis was monitored in cells growing on M63 minimal medium supplemented with ^35^S-labelled methionine and cysteine. At the indicated time points, 100 µL culture were directly precipitated with TCA and separated by SDS-PAGE. Radioactively labelled proteins were visualized by autoradiography.

The partial dissipation of the pmf should reduce the metabolic activity of the YohP-producing cells and this was monitored in a metabolic labeling experiment. Cells were grown on minimal media in the presence of ^35^S-labeled methionine and cysteine to monitor the global protein synthesis rate. This demonstrated that the protein synthesis rate of *yohP*-expressing cells was reduced in comparison to wild type cells (**Fig. 5C**).

### YohP production is linked to the stringent response

One striking effect of YohP production is the down-regulation of several enzymes involved in pyrimidine biosynthesis. Due to the high energy demand of their *de-novo* synthesis, purine and pyrimidine biosynthesis in *E. coli* are strictly regulated (65,66). One mechanism that coordinates nucleotide biosynthesis with nutrient availability in bacteria is the stringent response (8,67–69). The stringent response describes the accumulation of the stress-signaling molecules pppGpp and ppGpp, which induce multi-layered changes in cellular metabolism, including changes in transcription, translation and protein targeting (30,31,70,71). For analyzing whether YohP production influences the stringent response, the (p)ppGpp levels in cell extracts were determined by capillary electrophoresis combined with mass spectrometry (CE-MS) (72,73) (**Fig. 6A**). This approach revealed 4-fold increased ppGpp levels in the YohP-producing strain compared to the wild type strain. The pppGpp levels in all three strains were significantly lower than the ppGpp levels, which is in line with published data (74) and mainly caused by GppA, which converts pppGpp into ppGpp (75). The Δ*yohP* strain showed a small increase of the pppGpp levels, but whether this is the result of the reduced TnaA and/or indole levels needs to be further analyzed. Thus, the induction of *yohP* causes an up-regulation of ppGpp, which in turn could repress pyrimidine biosynthesis. This was further validated by expressing pRS1-YohP in a strain that lacks RelA, the key enzyme for (p)ppGpp production in *E. coli* (8). As a read-out for enzymes of the pyrimidine biosynthesis, we monitored the levels of the dihydroorotate dehydrogenase PyrD. The PyrD levels in strain MG1665 containing pRS1-yohP were reduced in comparison to the MG1665 wild type, confirming the results from the mass spectrometry. Importantly, there was no significant difference in the PyrD levels between the Δ*relA* strain and the Δ*relA* strain containing pRS1-*yohP* (**Fig. 6B**), which is explained by the reduced amounts of (p)ppGpp that are synthesized when RelA is missing (76). The inner membrane protein YidC served as loading control in these experiments. In conclusion, the down-regulation of enzymes involved in pyrimidine biosynthesis is likely the consequence of the stringent response.

**Fig. 6.**
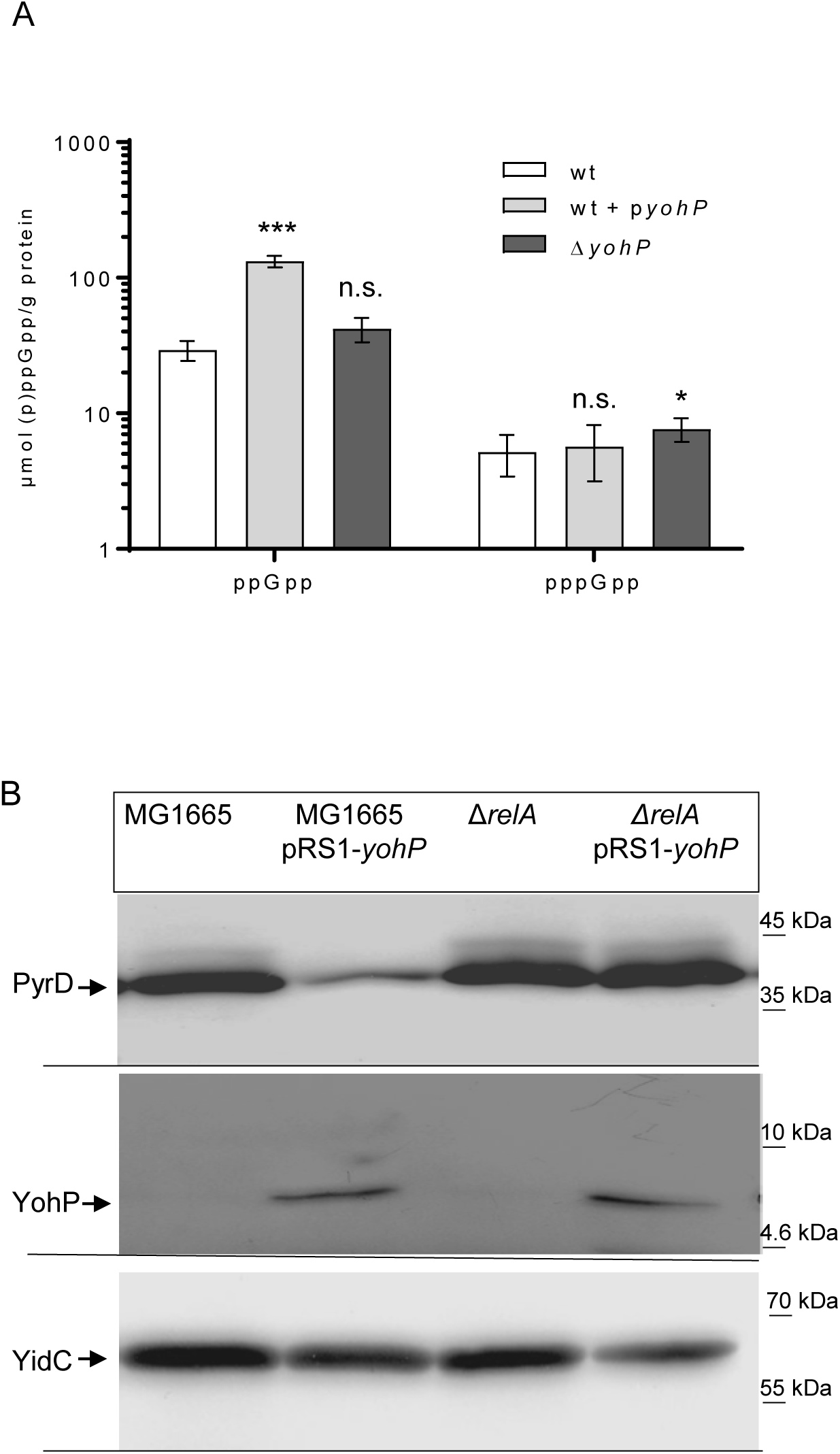
YohP production induces the stringent response. **A**. *E. coli* cells were grown on LB medium up to OD_600_ = 0.8 and *yohP* expression was induced at OD_600_ = 0.3. Subsequently, 20 x 10^8^ cells were lysed and extracted as described in Material and Methods. Samples were spiked with heavy [^15^N] (p)ppGpp standards before extraction. (p)ppGpp levels in the extracted samples were determined by MS-coupled capillary electrophoresis. The assay was performed with three biological replicates and three technical replicates each (n = 9). **B**. The *E. coli* strains were grown as in A and 2 x 10^8^ cells were TCA precipitated and separated by SDS-PAGE, followed by immune detection using α-PyrD, α-Flag and α-YidC antibodies. To determine the significance of the results, the P-value was calculated using the ‘unpaired, two-tailed t-test’ of the program Graph Pad PRISM 6 or the one-way Anova analyses and Turkey’s honest significance test using the values in the wild type strain as reference. The P-values are depicted as asterisks (*) above the graphs as following: n.s. = P > 0,05; * = P ≤ 0,05; ** = P ≤ 0,01.

## Discussion

Although technical improvements have dramatically advanced the identification of SMPs in bacterial proteomes (77), their functional characterization remains a bottle-neck for understanding their role in bacterial physiology (78–83). In the current study we employed a multidisciplinary approach to investigate and gain new insights on the function of YohP in *E. coli.* Our data demonstrate that YohP insertion into the *E. coli* membrane causes membrane damage that leads to the dissipation of the proton motive force. This in turn stimulates the stringent response and induces significant changes in the proteome.

One important observation of our study is that the deletion of *yohP* does not cause any detectable phenotype under the conditions tested and that it has only a minor effect on the *E. coli* proteome. Only TnaA showed a significant down-regulation in the absence of YohP, and although indole, the product of the TnaA reaction, has been linked to stress signaling and resistance in bacteria (56,57,84), we only observed slightly reduced indole production in the *E. coli* Δ*yohP* strain. This likely reflect the fact that TnaA was still detectable by immune detection in the Δ*yohP* strain. Besides TnaA, only YgeY showed an obvious down-regulation upon *yohP* deletion. YgeY is putative peptidase that was identified in a genetic screen for mutants defective in cell envelope biogenesis (46). STRING analyses (55) also link it to the salvage pathway of purines, but its exact function is still unknown.

Thus, although our data show that YohP production is up-regulated when cells enter stationary phase, transition into and survival during stationary phase is not strongly compromised in the absence of YohP. This could be due to some functional redundancies between the abundant SMPs in *E. coli* (12,14,26,77).

Contrary to the minor proteomic effects of the Δ*yohP* strain, YohP overproduction results in major proteomic rearrangements. The highest up-regulation was observed for proteins that sense membrane damage, such as PspA, PspC and GlpC (48,85). PspC together with the transcriptional activator PspF activates the phage-shock response, which stabilizes the bacterial membrane and protects it against envelope stress (85–87). PspA is suggested to sense proton leakage through damaged membranes and to reduce further leakage by forming a carpet-like seal below the membrane (86,88). The upregulation of the phage-shock response is in line with the impaired membrane potential and the observed changes in membrane composition and fluidity in response to YohP insertion. PspF also activates the alternative sigma factor σ^54^ (RpoN) (89), which responds to nitrogen starvation and regulates more than 130 genes involved in nitrogen metabolism, the stringent response and stress signaling (90,91). This potentially explains the increased production of the alarmones (p)ppGpp upon YohP insertion. RelA-dependent (p)ppGpp production is typically induced by uncharged tRNAs as a consequence of amino acid starvation (92,93). Amino acid uptake in *E. coli* involves secondary transporters that use the electrochemical gradient as energy source, but their activity is compromised when the membrane potential is dissipated (94). Amino acid starvation upon YohP-induced membrane damage will further stimulate the stringent response and also explains the differential expression of many enzymes involved in amino acid metabolism that we observe in our study. A major target of (p)ppGpp is the machinery required for ribosome biogenesis and translation (8,95), which rationalizes the reduced global protein synthesis when YohP is expressed. However, our proteome analysis did not show any indication for reduced ribosomal protein levels, with the exception of RpsU (bS21), which is required for translation initiation and transcriptionally inhibited by ppGpp (96). But in general, the (p)ppGpp levels upon YohP production apparently do not reach levels that are sufficient to completely inhibit ribosome biogenesis (8,43,95).

Several enzymes involved in the assembly of the outer membrane and the peptidoglycan biosynthesis are also upregulated upon YohP production and the upregulation of some of these proteins *(e.g.* MlaB, LptE, MepM) has been linked to increased resistance phenotypes (97–100). Thus, it is tempting to speculate that the YohP-linked changes in the cell envelope promote the overall survival under stress conditions.

Many of the down-regulated proteins are linked to amino acid, ribose-phosphate and nucleotide metabolism, which are all coordinated by the stringent response (65,101). The inhibition of purine nucleotide biosynthesis by (p)ppGpp has been reported before (68,102) and is also visible in our study. In contrast, less is known about a possible impact of (p)ppGpp on pyrimidine metabolism. However, enzymes involved in pyrimidine biosynthesis were identified as potential (p)ppGpp targets in ppGpp-capture studies (71,102). Furthermore, transcription of *pyrD* was shown to be strongly inhibited in the presence of ppGpp (103), which explains why the PyrD levels are reduced upon YohP-induced ppGpp accumulation. Thus, the induction of the stringent response as a consequence of YohP-induced membrane damage leads to an overall reduction of the nucleotide pool.

Many of the effects seen upon YohP insertion into the membrane are detrimental to the cell, which explains the reduced growth rate. It is important to emphasize that endogenous YohP levels are quite high during stationary phase and match the plasmid-encoded levels (*c.f.* Fig. 1). The major difference between the endogenous YohP production and the experimental system used here is that our system monitored the effect of premature YohP production in exponentially growing cells. This still raises the question about the advantage of producing an apparently inhibitory protein when *E. coli* enters stationary phase. It could be related to the increased stress resistance of metabolically silent cells, which cumulates in so-called persister cells, which can escape the effects of many antibiotics by entering a state of metabolic dormancy (39,104,105). Nonheritable antibiotic resistance is also facilitated by a reduced pmf (106) and the purpose of producing YohP during stationary phase could be to reduce metabolic activity as a trade-off for increased resistance against non-favorable conditions. Reducing metabolic activity is a common strategy of bacteria during stationary phase and well exemplified by ribosome hibernation factors which transiently inactive ribosomes and reduce global protein synthesis (43,64,107).

In conclusion, our data indicate that the primary effect of YohP production in *E. coli* is to induce a non-lethal membrane damage, which in turn leads to a partial metabolic silencing that ultimately protects cells against further damage and helps them to save cellular resources when nutrients are scarce.

### Experimental Procedures

#### Strains and plasmids

The *E. coli* K-12 strain MG1665 (108) was used as wild type strain and the MG1665 variant containing the chromosomally tagged *yohP* copy (GSO333) was a gift from Dr. Gisela Storz, National Institute of Health, USA. The Δ*relA* and Δ*tnaA* strains were obtained from the KEIO collection (109) via Thermo Fisher Scientific (Darmstadt, Germany). *E. coli* cells were routinely grown on LB-medium at 37° C in either liquid media or on solid media (LB medium + 1.5% agar-agar) unless otherwise stated. The plasmid pRS1-YohP has been described previously (29) and the *yohP*-coding sequence including the His_6_ sequence of pRS1-YohP was transferred via Gibson-assembly into pBad24 (110) to generate pBad24-YohP. The His6-tag in pRS1-YohP was replaced by a Flag3-tag via Gibson assembly. All nucleotide primer used in this study are listed in Supplementary **Table S3**. All PCR reactions were carried out using Q5^®^ High-Fidelity DNA Polymerase (M0491) and Phusion^®^ High-Fidelity DNA Polymerase (NEB Biolabs) according to manufacturer’s instructions. The Gibson Assembly^®^ Protocol from New England Biolabs was used for incorporating DNA sequences into the plasmids used in this study – according to manufacturer’s instructions.

For generating the Δ*yohP* strain, the *E. coli* K12 MG1655 strain was transformed with the temperature sensitive pKD46 plasmid (41) and cells were grown at 30 [. The chloramphenicol acetyltransferase gene flanked by FRT sites was amplified from plasmid pKD3. Cells with the pKD46 plasmid were grown in LB medium at 30[and induced with 10mM L-arabinose after OD_600_ reached approximately 0,1 to induce expression of the λ-red genes The growth was continued at 30[until OD_600_=0,4. Then aliquots of 1ml were taken into two 1,5 mL centrifuge tubes and kept in ice for 10 minutes, centrifuged for 10 minutes at 4000 x g and 4 °C after which the pellet was resuspended in 1ml of ice-cold sterile H_2_O. The linear PCR fragment containing the chloramphenicol acetyltransferase gene flanked by FRT sites and sequences adjacent to the *yohP* gene was then electroporated into the MG1655 pKD46 strain followed by incubation for 2 h with moderate shaking at 30 °C. Aliquots were plated on Cm-containing plates and Δ*yohP* colonies were validated via colony PCR.

### SDS-polyacrylamide gel electrophoresis (SDS-PAGE) and western blot analyses

Proteins were separated via denaturing SDS-polyacrylamide gel electrophoresis (SDS-PAGE). Two different gel systems were used in this study: 15% Tris-Glycine PAGE for separation of proteins between 10-100 kDa and 16.5% Tris-Tricine PAGE for the separation of small proteins between 0,5-10 kDa (29). Samples were denatured at 37 °C or 56 °C for 10 min in Laemmli loading buffer (278 mM Tris-HCl, pH 6.8, 44.4% glycerol, 4.4% SDS, 0.02% bromophenol blue) containing fresh DTT at a final concentration of 25 mM. For the immune detection of proteins in whole cells, routinely 1 x 10^8^ cells were precipitated with 5 % trichloroacetic acid (TCA, final concentration) and incubated for 30 min on ice. The sample was subsequently centrifuged at 30.000 x g and 4 °C for 15 min, the pellet was resuspended in 20 µl of SDS loading buffer and incubated for 10 min at 56 °C before loading on SDS-PAGE. The SDS-PAGE-separated proteins were then transferred via tank blot onto 0.45 µm nitrocellulose membranes (GE Healthcare, Freiburg, Germany) at 750 mA for 2 hours. For the detection of YohP, the gel was transferred by semi dry western blotting onto 0,22µm PVDF PsQ membranes (Merck, Darmstadt, Germany). Membranes were blocked with 5% milk powder in T-TBS buffer for at least 1 h before the addition of the primary antibodies.

α-Flag antibodies (mouse α-Flag M2) were obtained from Merck (Darmstadt, Germany). PyrD antibodies (rabbit α-PyrD, CSB-PA812570XA01FQR) and PspC antibodies (rabbit α-PspC, CSB-PA365328XA01SZB) were obtained from Cusabio via Biozol (Hamburg, Germany). PspA antibodies were a kind gift from Dirk Schneider, Univ. Mainz. Peptide antibodies against RpoS have been described before (111). Antibodies against YidC were raised against the purified proteins and have been reported before (112,113). Polyclonal TnaA antibodies were obtained from AssayPro (No. 33517-05111; St. Charles, USA) and α-His antibodies were from Invitrogen (MA1-21315). A horseradish peroxidase-coupled secondary antibody from Caltag Laboratories (Burlingam, CA, USA) was used for detection; blots were incubated for 1 min with home-made ECL reagent and signals were detected by a CCD camera. Western blot samples were analyzed by using *ImageQuant* (GE Healthcare) or the *ImageJ/ Fiji* plug-in software (NIH, Bethesda, USA). All experiments were performed at least twice as independent biological replicates and representative gels/blots/images are shown. When data were quantified, at least three independent biological replicates with several technical replicates were performed and the signal intensity observed for wild type cells or cell extracts was set to 100%.

### Mass spectrometry of *E. coli* cells

*E. coli* MG1655, *E. coli* MG1655 pRS1-*yohP* plasmid and *E. coli* MG1655 Δ*yohP* were grown on phosphate rich medium (INV medium; 10 g yeast extract, 10 g tryptone/peptone, 1% glucose, 1mM KH_2_PO_4_, 166 mM K_2_HPO_4_) YohP producing cells were induced at the OD_600_ = 0,5 with 1mM IPTG and grown for additional 2h. All samples were harvested at the OD_600_ = 1,5-1,7 and subjected to cell lysis via 3 cycles through the Emulsiflex cell lysis system at 800 psi. The homogenate was centrifuged at 15.000 rpm in an SS34 rotor (Beckmann-Coulter, Krefeld, Germany) and the protein concentration in the supernatants were determined via the Pierce™ BCA Protein Assay Kit (Thermo Fisher, Freiburg, Germany).

A volume of cell lysate containing 200 µg of proteins was mixed 1:1 with SDS buffer (10% SDS, 100 mM tetraethyl ammonium bicarbonate (TEAB) pH 8.5) and subjected to tryptic digestion following the S-Trap protocol (114) with small modifications. Briefly, proteins were reduced by addition of 4 µL TCEP for 30 min at 37 °C with mild agitation, and alkylated with 8 µL iodoacetamide (IAA) for 30 min in the dark with mild agitation. Samples were acidified with 12 µL 12% phosphoric acid, diluted with 750 µL of the S-Trap binding buffer (1 M TEAB Buffer and Methanol (10:90)) and transferred to S-Trap mini columns (PROTIFI, Fairport, USA). Three washing cycles with 400 µl S-Trap binding buffer were done prior to digestion for SDS removal. MS-grade Trypsin (Serva, Heidelberg, Germany) was added in an enzyme to protein ratio 1:50 using 125 µL digestion buffer (50mM ammonium bicarbonate (ABC), pH 8.5) as media and the column was incubated overnight at 37 °C with light agitation. Peptides were eluted stepwise in 80 µL 50mM ABC, 0.2% formic acid, 50% acetonitrile (ACN). The pooled fractions were diluted 1:1 with 0.1% trifluoroacetic acid (TFA) and desalted with C18-SPE cartridges (25 mg, Biotage, Uppsala, Sweden). After equilibration with 2 mL ACN, 1 mL 50% ACN/1% acetic acid and 2 mL 0.1% TFA the samples were loaded onto the cartridge, washed with 2 mL 0.1% TFA and eluted with 1 mL 80% ACN/0.1% TFA. The eluted fractions were dried using an Eppendorf concentrator (Eppendorf) and stored at -20 °C.

Dried peptides were reconstituted in 5% ACN with 0.1% formic acid (FA). Peptides were loaded onto an Acclaim PepMap C18 capillary trapping column (particle size 3 µm, L = 20 mm) and separated on a ReproSil C18-PepSep analytical column (particle size = 1.9 µm, ID = 75 µm, L = 50 cm, Bruker Corporation, Billerica, USA) using a nano-HPLC (Dionex U3000 RSLCnano) at a temperature of 55 °C. Trapping was carried out for 6 min with a flow rate of 6 μL/min using a loading buffer composed of 0.05% trifluoroacetic acid in H2O. Peptides were separated by a gradient of water (buffer A: 100% H_2_O and 0.1% FA) and acetonitrile (buffer B: 80% ACN, 20% H_2_O, and 0.1% FA) with a constant flow rate of 250 nL/min. The gradient went from 4% to 458% buffer B in 9120 min. All solvents were LC-MS grade and purchased from Riedel-de Häen/Honeywell (Seelze, Germany). Eluting peptides were analyzed in data-independent acquisition mode on an Orbitrap Eclipse mass spectrometer (Thermo Fisher Scientific) coupled to the nano-HPLC by a Nano Flex ESI source.

MS1 survey scans were acquired over a scan range of 350 – 1650 m/z in the Orbitrap detector (resolution = 120k, automatic gain control (AGC) = 5e^5^, and maximum injection time = 100 ms). Sequence information was acquired by a tMS2 method with the following settings. MS2 scans were generated in a scan range of 350 – 2000 m/z with isolation windows proposed by Muntel et al. (115). Fragmentation was induced by higher-energy collisional dissociation (HCD) at 27% normalized collision energy (NCE). For Orbitrap MS2, an automatic gain control (AGC) of 1xe^6^ with dynamic injection time and a minimum of 6 desired points across the peak (resolution = 30k).

MS raw files were processed with the open-source software DIA-NN (version 1.8.1) (116) using a library-free approach. The predicted library was generated by in silico digesting the *Escherichia coli* UniProt reference proteome (UP000000625, 4399 entries, downloaded on March 2023) with Trypsin/P. Deep learning-based spectra- and RT-prediction was enabled. Peptide length range was set to 6 - 35 amino acids, missed cleavages to 2 and precursor charges to 1-5. Methionine oxidation and N-terminus acetylation were set as variable modifications, and cysteine carbamidomethylation as a fixed modification. The maximum number of variable modifications per peptide was limited to 1. MS1 and MS2 accuracies were automatically calculated by DIA-NN. Protein inference was performed using genes with the heuristic protein inference option enabled. The neural network classifier was set to single-pass mode and the quantification strategy was selected as ‘QuantUMS (high precision)’. Reported data from DIA-NN were analyzed using the MS-DAP-R package (45) with the following parameters: filter minimum detect = 2, filter minimum quant = 2, filter minimum peptide per protein = 1, filter by contrast = true, normalization algorithm = vsn followed by mode between_protein, differential expression analysis algorithm = deqms and q-Value threshold = 1%.

### Downstream analysis of proteomics data

Outputs from the differential expression analysis were imported in RStudio for downstream analysis, and all plots were generated using ggplot2. For all downstream analysis, proteins with a statistically significant differential expression (q-value < 0.05) and with a log2-fold change of at least |1| were selected.

GO enrichment analysis was performed using clusterProfiler (117) with the org.EcK12.eg.db as source for annotations. The obtained p-values in the GO analysis were corrected using the Benjamin-Hochberg method with a 5% confidence level. GO analysis was performed separately for up- and downregulated proteins, and for GO annotations linked to biological processes and cell compartment.

Lists of differentially expressed proteins were also uploaded to STRING (55) for network analysis. Interaction networks were generated for up- and downregulated proteins separately and combined. Non-interacting nodes were removed from the final output, and a minimum interaction score of 0.4 was used.

### Indole measurements

*E. coli* cells for indole measurements were grown on INV medium or tryptone broth (10g/L tryptone, 5g NaCl/L) up to the indicated optical density. Cells containing pBad-YohP were induced with 0.1 % arabinose. After centrifugation at 20.000 g for 30 min at 4 °C, the indole concentration in the supernatant was determined using the Kovacs regent (dimethyl-amino-benzaldehyd) (Roth, Karlsruhe, Germany) (118). In brief, 10 µl of the supernatant were incubated with 200 µl Kovacs reagent in a 96-well plate for 1 h at room temperature. The formed cyan-dye rosindole was determined by measuring the absorption at 571 nm with a Tecan Spark plate reader, using an indole standard curve (0-1 mM indole) as reference.

### Lipidomic analyses of inner membrane vesicles

Lipid composition was determined in inner membrane vesicles (INVs) of the indicated strains. Cells were grown on INV medium up to OD_600_ of 1.5, harvested and washed twice in buffer A (50 mM TeaOAc, pH 7.5; 250 mM sucrose, 1mM EDTA, 1 mM DTT). Cells were resuspended in 1 ml buffer A/g cell weight and lysed in a cooled French pressure cell. The lysate was then centrifuged at 150.000 g for 2 h in Beckmann Ti50.2 rotor and the pellet containing the membranes were resuspended in buffer A. These crude membranes were loaded on a discontinuous sucrose gradient (0.77 M; 1.44 M; 2.02 M sucrose in buffer A + 0.5 mM PMSF) and centrifuged in Sorvall SureSpin106.4 swing-out rotor at 25.000 rpm for 17h. After centrifugation, the inner membrane phase was collected, diluted with 50 mM TeaOAc, pH 7.5 and centrifuged for 2 h at 150.000 g in a Beckmann TI50.2 rotor. The pellet was resuspended in buffer A and samples were stored in small aliquots at -80 °C.

#### Lipid extraction for mass spectrometry lipidomics

Mass spectrometry-based lipid analysis was performed by Lipotype GmbH (Dresden, Germany) as described (119). Lipids were extracted using a two-step chloroform/methanol procedure (120). Samples were spiked with internal lipid standard mixture containing: cardiolipin 14:0/14:0/14:0/14:0 (CL), ceramide 18:1;2/17:0 (Cer), diacylglycerol 17:0/17:0 (DAG), hexosylceramide 18:1;2/12:0 (HexCer), lyso-phosphatidate 17:0 (LPA), lyso-phosphatidylcholine 12:0 (LPC), lyso-phosphatidylethanolamine 17:1 (LPE), lyso-phosphatidylglycerol 17:1 (LPG), lyso-phosphatidylinositol 17:1 (LPI), lyso-phosphatidylserine 17:1 (LPS), phosphatidate 17:0/17:0 (PA), phosphatidylcholine 17:0/17:0 (PC), phosphatidylethanolamine 17:0/17:0 (PE), phosphatidylglycerol 17:0/17:0 (PG), phosphatidylinositol 16:0/16:0 (PI), phosphatidylserine 17:0/17:0 (PS), cholesterolester 20:0 (CE), sphingomyelin 18:1;2/12:0;0 (SM), triacylglycerol 17:0/17:0/17:0 (TAG). After extraction, the organic phase was transferred to an infusion plate and dried in a speed vacuum concentrator. First step dry extract was resuspended in 7.5 mM ammonium acetate in chloroform/methanol/propanol (1:2:4, V:V:V) and second step dry extract in 33% ethanol solution of methylamine in chloroform/methanol (0.003:5:1; V:V:V). All liquid handling steps were performed using Hamilton Robotics STARlet robotic platform with the Anti Droplet Control feature for organic solvents pipetting.

#### MS data acquisition

Samples were analyzed by direct infusion on a QExactive mass spectrometer (Thermo Scientific) equipped with a TriVersa NanoMate ion source (Advion Biosciences). Samples were analyzed in both positive and negative ion modes with a resolution of Rm/z=200=280000 for MS and Rm/z=200=17500 for MSMS experiments, in a single acquisition. MSMS was triggered by an inclusion list encompassing corresponding MS mass ranges scanned in 1 Da increments (121). Both MS and MSMS data were combined to monitor CE, DAG and TAG ions as ammonium adducts; PC, PC O-, as acetate adducts; and CL, PA, PE, PE O-, PG, PI and PS as deprotonated anions. MS only was used to monitor LPA, LPE, LPE O-, LPI and LPS as deprotonated anions; Cer, HexCer, SM, LPC and LPC O-as acetate adducts.

#### Data analysis and post-processing

Data were analyzed with in-house developed lipid identification software based on LipidXplorer (122). Data post-processing and normalization were performed using an in-house developed data management system. Only lipid identifications with a signal-to-noise ratio >5, and a signal intensity 5-fold higher than in corresponding blank samples were considered for further data analysis.

### Time-resolved DPH anisotropy analysis of inner membrane vesicles (INVs)

Inner membrane vesicles were prepared as described above and lipid concentrations of the INV dispersion was determined by the Bartlett assay (123). The homogeneity of the INV samples was validated by dynamic light scattering (DLS) measurements, which were conducted at 20°C using a Nano-ZS Zetasizer from Malvern Panalytical (Kassel, Germany). The instrument was equipped with a 633 nm He-Ne laser and detected at an angle of 173°. Data were collected and analyzed using Zetasizer software (version 7,13), which also calculated the viscosity and refractive index of the medium from its database. The software automatically optimized the attenuator and measurement position and was also used to determine particle size and size distribution. The INV samples were adjusted to a total protein concentration of 1 mg and 0.1 mol% DPH (< 1 vol% DMSO) along with the buffer solution (10 mM Tris, 100 mM NaCl, 0.02% NaN3, pH 7,4 at 20°C) were added. The samples were incubated for 10 minutes at 20°C with stirring at 400 rpm. Time-resolved anisotropy measurements were performed at 20 °C using a FluoTime 300 spectrometer from PicoQuant (Berlin, Germany) with 100 µL volume Ultra-Micro cells from Hellma (Mulheim, Germany) having an optical path of 10 x 2 mm. The setup and initial data analysis were conducted using EasyTau Software (version 2.2.3293). Excitation was done at 355 nm with a laser polarization of 0° at a frequency of 16,67 MHz, laser intensity of 7,2, and a pulse width of 25 ps. Emission was measured through a 355 nm long-pass filter at 430 nm with a 5 nm detection band pass, and the emission polarizer set to 0°, 54,7°, and 90°, respectively. The G-factor was recorded with the same setup with laser polarization at 90° and calculated by manually aligning emission decays at polarizations of 0° and 90°. The anisotropy decay was obtained by a tail fit of the decay curves with vertical and horizontal polarizer and fitted assuming a mono-exponential decay in terms of the initial anisotropy, the correlation time of the angular motion of DPH, and the limiting anisotropy, r_∞_, extrapolated to infinite time after excitation. The latter parameter represents the angular constraints to the tumbling of the probe so that a large r_∞_ stands for relatively high order and tight packing of the acyl chains. The goodness of fit was assessed using reduced chi-square values and bootstrap error analysis, although these are not shown. The time-resolved DPH anisotropy analysis was performed using three independent biological replicates of INVs, each containing three technical replicates.

### Membrane potential measurement in whole cells

The *E. coli* K12 MG1655 strain with the indicated plasmids were grown on LB and protein production was induced at OD600=0.5 and further incubated at 37 [for 2 h. Aliquots of 1×10^9^ cells were collected from each sample and washed once with sterile PBS. When indicated 1mM of CCCP (100mM stock in DMSO) was added. All samples were adjusted to a total volume of 100 µl of and further incubated for 30 minutes at 37 [. The samples were washed with PBS and diluted with 500 µl of PBS supplemented with 2µg final concentration of the dye DiBAC_4_(3) (Stock solution: 10 mg/ml in DMSO, working solution: 200 µg/ml in sterile deionized H_2_O, freshly prepared). Samples were incubated at room temperature for 20 minutes in the dark, centrifuged at 13.000 rpm for 3min at room temperature, washed 2x with sterile PBS, and incubated with 2% paraformaldehyde in a total volume 100 µl PBS for 10 min at room temperature. Samples were then washed with PBS and finally resuspended in 200 µl PBS. 100 µl from this solution was transferred on a black 96 well plate (Greiner, Frickenhausen Germany) and fluorescence was measured in a Tecan Spark plate reader after excitation at 485 nm and an emission of 510 nm.

### *In vivo* metabolic labeling

*E .coli* MG1655 containing either plasmid pRS1 or pRS1-yohP were grown and induced for 2 h on M63 minimal medium (124) supplemented with 20 amino acids. Cells were harvested and washed 3 times with M63 medium lacking methionine and cysteine, resuspended in same medium and induced for additional 30 minutes. An aliquot of 2×10^8^ cells were then added to 1m M63 medium and supplemented with 1 μCi ^35^S-methionine and cysteine labeling mix (Perkin Elmer, Rodgau, Germany). 100 microliters samples were taken from each culture at different time points after the addition of radioactive labeling mix and directly precipitated with 5% TCA, separated with SDS PAGE and visualized using autoradiography.

### Viable cell staining and counting

The assay was performed using the QUANTOM Tx^TM^ Microbial Cell Counter and the QUANTOM^TM^ Viable Cell Staining Kit obtained from BioCat GmbH (Heidelberg, Germany). The kit stains live bacterial cells to be counted. The optical density of the cell-culture was determined and an aliquot corresponding to approx. 1 x 10^8^ cells were collected. The cells were washed with PBS and the culture media was completely removed. Cells were resuspended in PBS buffer and incubated with different CCCP concentrations or DMSO as a control for 1 h at 37 °C. Subsequently, the cells were collected by centrifugation and washed with QUANTOM Cell Dilution Buffer, centrifuged and resuspended in 80 µL QUANTOM Cell Dilution Buffer. 10 µL of this suspension were transferred into a fresh tube and incubated with 2 µL of the QUANTOM^TM^ Viable Cell Staining Dye was added and mixed gently and carefully. The cells were then incubated at 37°C for 30 minutes in the dark. Thereafter 8.0µL QUANTOM^TM^ Cell Loading Buffer I was added and mixed gently without creating bubbles. 5.0µL of this mixture was loaded onto a QUANTOM^TM^ M50 Cell Counting Slide and centrifuged at 300 x g for 10 minutes in a QUANTOM^TM^ Centrifuge at room-temperature. The slide was then inserted into the QUANTOM Tx^TM^ cell counter and cells were counted with the light intensity level set to either 7 or 9. The obtained viable cell numbers per ml were then blotted against the CCCP concentration.

### Extraction and determination of (p)ppGpp by capillary electrophoresis-mass spectrometry (CE-MS)

*E. coli* cells were grown on LB medium up to OD_600_ = 0.8 and 20 x 10^8^ cells were lysed with pre-chilled formic acid (final concentration 1 M) and stored at -80 °C until further extraction. Samples were thawed and spiked with heavy [^15^N]_5_ (p)ppGpp standards before extraction. Following incubation for 30 minutes with vortexing every two minutes, samples were diluted with 50 mM NH_4_OAc, pH 5.5 and centrifuged (10 min, 3220 g, 4 °C). The supernatant was subjected to weak anion solid phase extraction using a GX-241 ASPEC system (Gilson Inc, Middelton, USA) and EVOLUTE® EXPRESS WAX cartridges (100 mg/3 ml (Tabless)) from biotage (Uppsala, Sweden), which were equilibrated with MeOH (1ml) and NH_4_OAc (1 ml, 50 mM, pH 4.5) before samples were loaded onto the column. Cartridges and analytes were washed with NH_4_OAc (1 ml, 50 mM, pH 4.5) and MeOH (1 ml). Samples were eluted using a MeOH:ddH_2_O:NH_4_OH buffer (20:70:10, 2 × 750 µl), diluted with ddH_2_O (2 ml) and lyophilized overnight. The precipitate was dissolved in in ddH_2_O (100 µl) and the solvent was removed using an Eppendorf vacuum concentrator for 5h at room temperature. The obtained precipitate was dissolved in 60 µl ddH_2_O. The extracted samples were stored at -20 °C until measurement. Sample preparation for the CE-MS measurement was performed by centrifugation of the samples and further dilution of the aqueous sample in a ratio of 1:1 with ddH_2_O.

Capillary electrophoresis (CE) was performed using an Agilent 7100 CE system (Agilent Technologies, Waldbronn., Germany). The CE was coupled to an Agilent G6495C Agilent QQQ mass spectrometer via a commercial Agilent jet stream (AJS) electrospray ionization (ESI) source and an Agilent liquid coaxial interface. The analyte solution was diluted with a ddH_2_O-isopropanol mixture (1:1 v-%), using an Agilent 1200 isocratic LC pump and a 1:100 splitter. The resulting flow was 10 µl/min. A bare fused silicia capillary (100 cm length, 50 μm internal diameter) was activated and flushed with 1 M NaOH and water for 10 minutes before the first measurement. In the beginning of each measurement, the capillary was washed with ddH_2_O (300 s) and ammonium acetate serving as separation buffer (35 mM, pH 9.75, 300 s). After sample injection with pressure of 100 mbar for 10 s and a buffer plug (50 mbar, 5 s), a voltage of +30 kV was applied for separation, which resulted in a stable current of 22 µA. The MS ran in negative ionization mode with a capillary voltage of - 2000 V and a nozzle voltage of 2000 V. The pressure RF was set to an upper limit of 90 V and a lower limit of 60 V. Nebulizer gas was set to 8 psi with a temperature of 150 °C and a flow of 11 l/min. The sheath gas had a temperature of 175 °C and a flow of 8 l/min. Peak assignment was performed with internal heavy standards and MS/MS transitions.

### Quantification and Statistical analysis

Western blot and autoradiography samples were analyzed by using the *ImageQuant* (GE Healthcare) or the *ImageJ/ Fiji* plug-in software (NIH, Bethesda, USA). All experiments were performed at least twice as independent biological replicates and representative gels/blots/images are shown. When data were quantified, at least three independent biological replicates with several technical replicates were performed. Mean values and standard deviations were determined by using either Excel (Microsoft Corp.) or GraphPad Prism (GraphPad Prism Corp. San Diego). For statistical analyses, a Student unpaired two-way t-test was performed.

## Data availability statement

All data are contained within the manuscript, with the exception of the mass spectrometry proteomics data, which have been deposited to the ProteomeXchange Consortium via the PRIDE (125) partner repository with the dataset identifier PXD063229.

## Supporting information

This article contains supporting information.

## Acknowledgements

We are grateful to Dr. Gisela Storz, NIH Bethesda, USA, for providing the *E. coli* GSO33 strain and to Dr. Dirk Schneider, Johannes Gutenberg University Mainz for providing the PspA antibody.

## Abbreviations and nomenclature

CCCP: cyanide-*m*-chlorophenyl hydrazine;
DiBAC_4_(3): Bis-(1,3-Dibutylbarbituric Acid) Trimethine Oxonol);
DPH: diphenylhexatriene;
GO: gene ontology;
IPTG: isopropyl β-D-1-thiogalactopyranoside;
INV: inner membrane vesicles;
SMP: small membrane protein

