## Supplemental information for "The small membrane protein YohP induces membrane depolarization and ppGpp accumulation in *Escherichia coli*"

**Table S1: Differentially expressed proteins in the Δ*yohP* strain (Table S1.exe).**

**Table S2: Differentially expressed proteins in the *yohP*-expressing strain (Table S2.exe).**

**Table S3. Oligonucleotide primers used in this study**

| Name | purpose | sequence (5´-3`) |
|---|---|---|
| pRS1-fw | pRS1 opening for Flag-tag cloning | taactcgagtagcataaccccttg |
| pRS1-rev | pRS1 opening for Flag-tag cloning | aaatatcatcttaaatacgccagtcacc |
| pRS1-Flagx3 | 3x Flag sequence attachment to yohP | gcgtatttaagatgatatttggcggcggcagcgactacaaggaccacgacgg-cgactacaaggaccacgacatcgactacaaggacgacgacgacaagtaactcgagtagcataaccc |
| pBad24-fw | pBad24 opening for Flag-tag cloning | taaatggtacccggggatcctctag |
| pBad24-rev | pBad24 opening for Flag-tag cloning | aaatatcatcttaaatacgccagtcacc |
| pBAd24-Flag3 | 3x Flag sequence attachment to yohP | gcgtatttaagatgatatttggcggcggcagcgactacaaggaccacgacgg-cgactacaaggaccacgacatcgactacaaggacgacgacgacaagtaaatggtacccggggatcc |
| CAT-fw | $Cm^R$ cartridge flanked by FRT sites | gtgtaggctggagctgcttc |
| CAT-rev | $Cm^R$ cartridge flanked by FRT sites | atgggaattagccatggtcc |
| YohP-Cm-fw | *yohP* indel | ggcttcgtttttcctatacttattcagcactcacaaataaaggaacgcc-agtgtaggctggagctgcttc |
| YohP-Cm-rev | *yohP* indel | ggaccatggctaattcccataattaattaatgtcatcaggtccgaaaata-acgagaatatttcagtctctc |
| YohP-control fw | control Δ*yohP* strain | cctatacttattcagcactcac |
| YohP-control rev | control Δ*yohP* strain | ctcgttattttcggacctgatgac |

:

**Supplementary Figure Legends:**

**Figure S1. Phenotypes of the Δ*yohP* and *yohP*-expressing strains. (A)** *E. coli* cells were grown on LB-medium and optical density was measured over time. (**B**) Precultures of the indicated strains were grown on LB medium up to $OD_{600}$=0.8, washed and serially diluted with PBS-buffer and spotted on MacConkey agar with maltose as single carbon source. (**C**) As in (B) but cells were spotted on LB plates with different pH values.

**Figure S2: Upregulated proteins in the *yohP*-induction strain are localized to the cell envelope and outer membrane.** Dotplots for the GO classification of Cellular Component (CC) enrichment analysis for the comparison between wild-type E. coli and the yohP-induction strain.

**Figure S3: YohP-producing cells contain well-connected groups of differentially expressed proteins.** String networks for the comparison between wild-type E. coli and the yohP-induction strain. Only differentially abundant proteins (q-value < 0.05 and fold-change > 2) were selected for the network analysis, and non-interacting nodes were removed. K-means clustering analysis of the network led to significant nodes ($p < 0.05$) grouped by different colors.

**Fig. S4: String network analyses of up- and down-regulated proteins upon YohP-production.** String analyses was performed as in Fig. S3 and up-regulated proteins are labeled in blue, while down-regulated proteins are labeled in red.

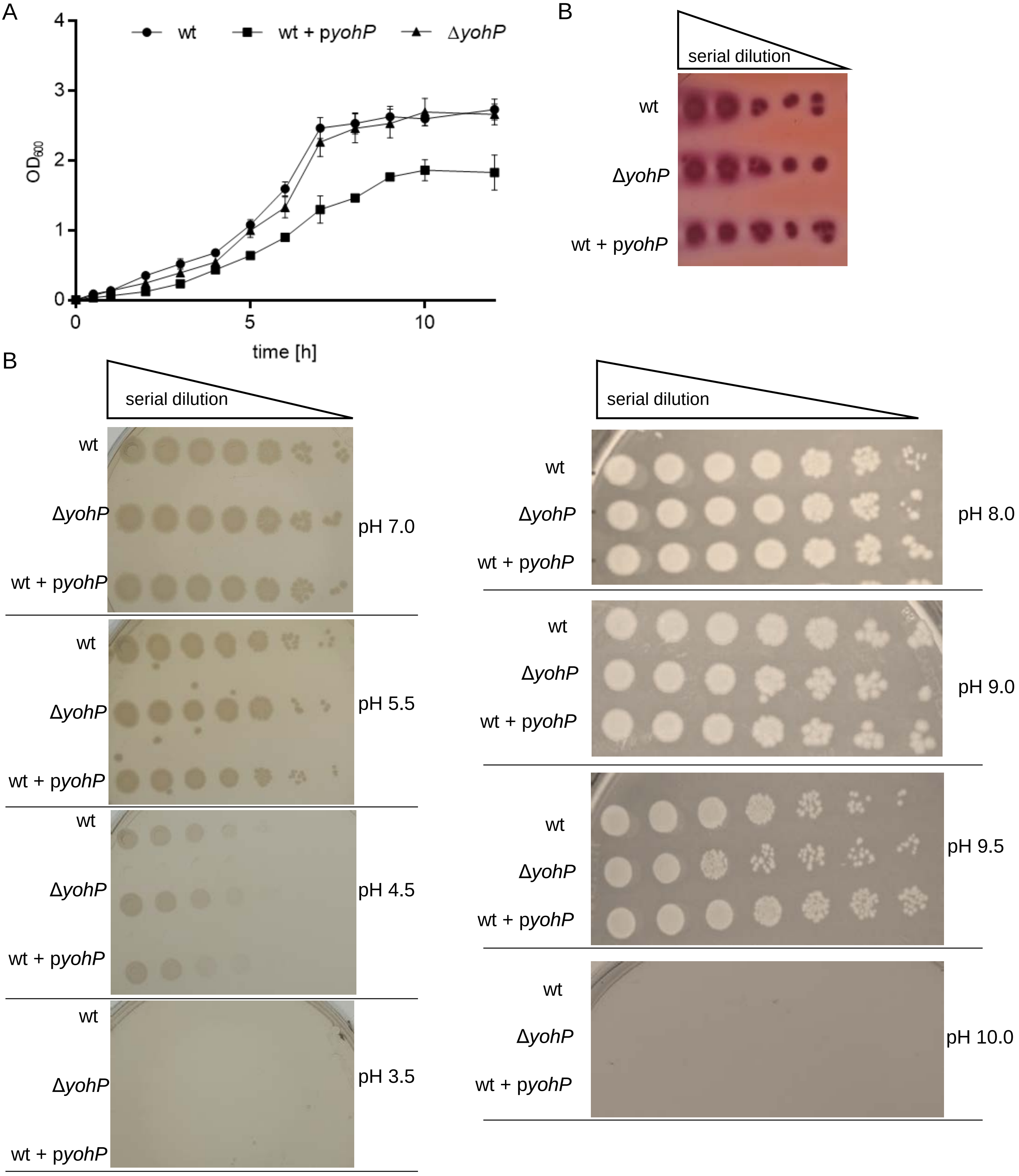


(Natriashvili et al., Fig. S1)

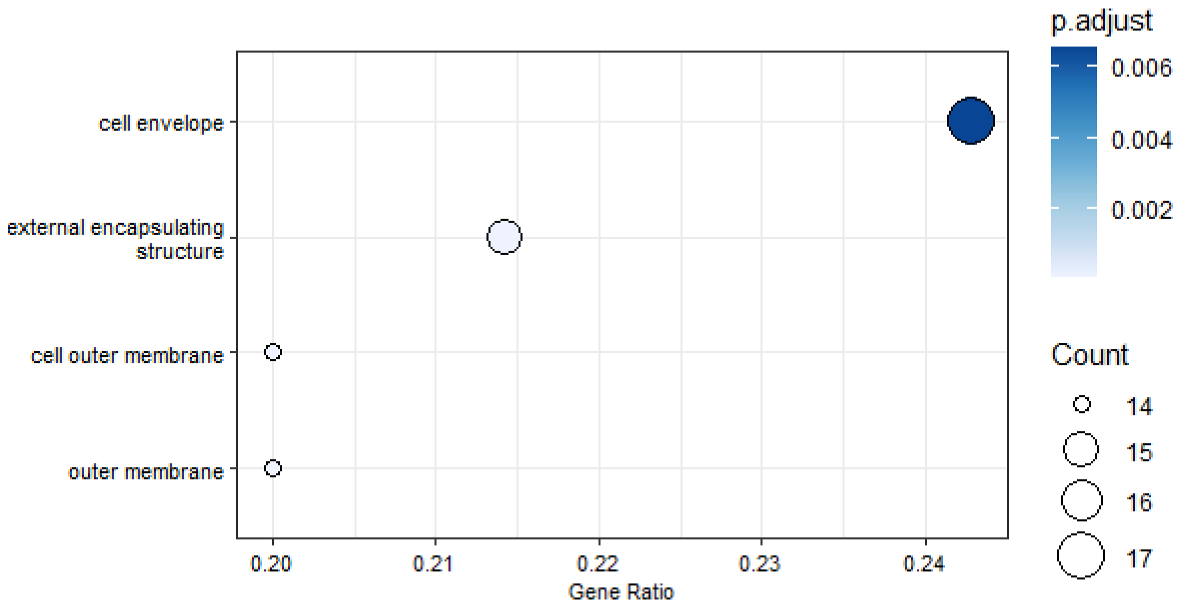


(Natriashvili et al., Fig. S2)

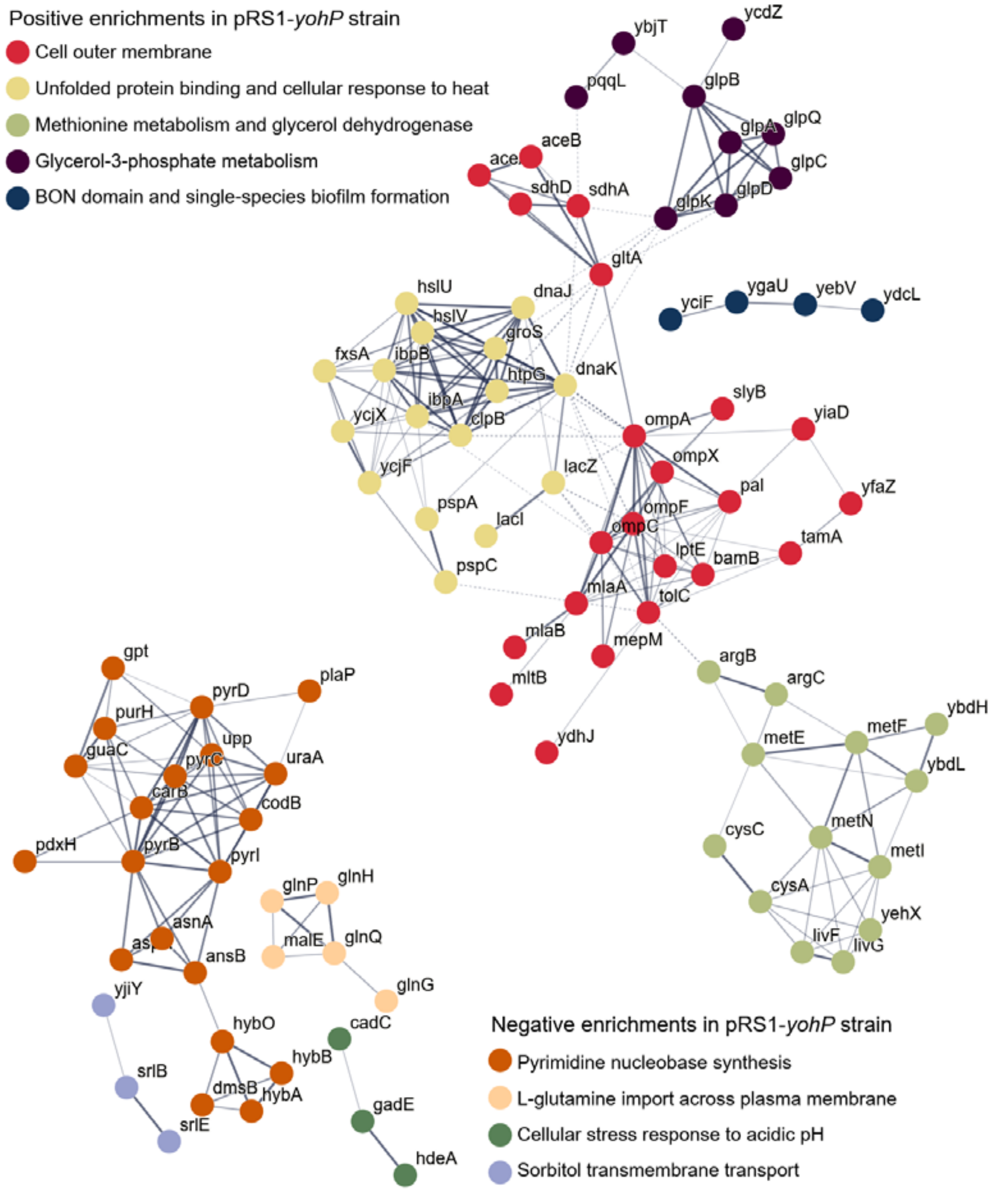


(Natriashvili et al., Fig. S3)

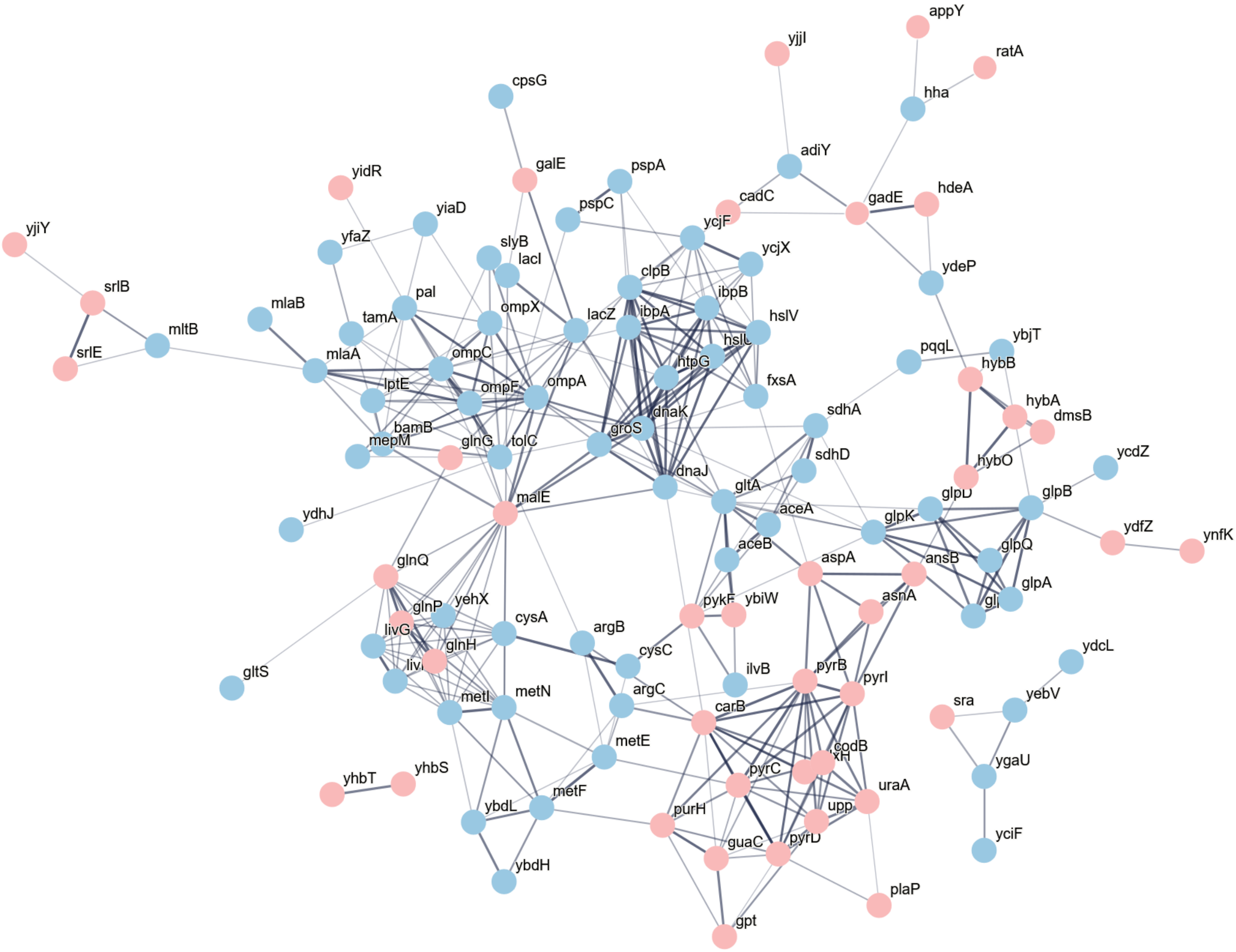


(Natriashvili et al., Fig. S4)
